# Osteogenic CpG Oligodeoxynucleotide, iSN40, Inhibits Osteoclastogenesis in a TLR9-Dependent Manner

**DOI:** 10.1101/2024.08.31.610656

**Authors:** Rena Ikeda, Chihaya Kimura, Yuma Nihashi, Koji Umezawa, Takeshi Shimosato, Tomohide Takaya

## Abstract

A CpG oligodeoxynucleotide (CpG-ODN), iSN40 (5’-GGA ACG ATC CTC AAG CTT-3’), was originally identified to promote osteoblast differentiation independent of Toll-like receptor 9 (TLR9). While CpG-ODNs are generally known to be recognized by TLR9 and inhibit osteoclastogenesis. This study investigated the anti-osteoclastogenic effect of iSN40. The murine monocyte/macrophage cell line RAW264.7 was treated with receptor activator of nuclear factor-κB ligand (RANKL) to induce osteoclast differentiation, and the effects of iSN40 on osteoclast formation were quantified by tartrate-resistant acid phosphatase (TRAP) staining and real-time RT-PCR. iSN40 completely inhibited RANKL-induced differentiation into TRAP^+^ multinucleated osteoclasts by suppressing osteoclastogenic genes (*Nfatc1, Ctsk*, and *Dcstamp*) and inducing anti-/non-osteoclastogenic genes (*Irf8, Adgre1*, and *Il1b*). Treatment with a TLR9 inhibitor, E6446, or mutation in the CpG motif of iSN40 abolished intracellular uptake and the anti-osteoclastogenic effect of iSN40. These results demonstrate that iSN40 is internalized subcellularly, recognized by TLR9 via its CpG motif, modulates RANKL-dependent osteoclastogenic gene expression, and ultimately inhibits osteoclast formation. Computational simulation of the iSN40 structure also suggested the importance of the superficial CpG motif for iSN40 function. Finally, iSN40 was confirmed to inhibit osteoclastogenesis of RAW264.7 cells cocultured with the murine osteoblast cell line MC3T3-E1, which is a model of bone remodeling. This study demonstrates that iSN40, which exerts both pro-osteogenic and anti-osteoclastogenic effects, may be a promising nucleic acid drug for osteoporosis.

## 1. Introduction

Bone homeostasis is continuously maintained by bone remodeling, which is accomplished by balancing two biological processes. One is bone formation, in which osteoblasts differentiate into osteocytes that form and mineralize extracellular matrix to produce bone tissue. The other is bone resorption, in which monocytes differentiate as pre-osteoclasts into multinucleated osteoclasts that degrade bone tissue. Osteogenesis and osteoclastogenesis regulate each other to maintain their balance [1], but bone remodeling becomes functionally uncoupled with age, leading to bone mass loss and eventually osteoporosis [2]. Today, the prevalence of osteoporosis worldwide is estimated to be 18.3% [3]. In the elderly, several factors such as sclerostin increase bone resorption relative to bone formation [4,5]. Therefore, a humanized monoclonal anti-sclerostin antibody, romosozumab, which induces osteogenesis and inhibits osteoclastogenesis [6], has been used in osteoporosis therapy [7]. The success of romosozumab demonstrates that a dual effect on both osteoblasts and osteoclasts is beneficial in the treatment of osteoporosis. However, antibody drugs are still expensive and difficult to mass produce for osteoporosis patients using current technology.

Oligonucleotides have been studied and applied in clinical settings as next-generation drugs due to their various advantages in chemical synthesis, variety of modifications, low-cost manufacturing, and storage stability [8]. For example, the single-stranded oligodeoxynucleotides (ODNs) with unmethylated CpG motifs (CG sequences), typically generated from viral and bacterial genomes, serve as Toll-like receptor 9 (TLR9) ligands and are used for anti-cancer therapy by modulating the immune system [9]. In parallel, we have reported that bacterial non-CpG-ODNs can become TLR9-independent aptamers that regulate cell proliferation, differentiation, and inflammation [10–15]. Such viral and microbial genome-derived ODNs are considered as promising seeds for nucleic acid drugs that exert bioactivities in various ways.

In the field of bone research, several osteogenetic ODNs (osteoDNs) have been identified that promote osteogenesis (Table 1). MT01, a non-CpG-ODN derived from the human mitochondrial genome, is the first osteoDN that induces osteogenic differentiation of the human osteoblast-like cell line MG63 [16,17], rat bone marrow mesenchymal stem cells [18], and the murine osteoblast cell line MC3T3-E1 [19]. Besides MT01, six designed CpG-ODNs (BW001, BW006, FC001, FC004, YW001, and YW002) were reported to promote the differentiation of MC3T3-E1 cells [19], but their TLR9 dependencies were not investigated. In contrast, CpG-2006 interferes with osteoblastic differentiation of human mesenchymal stem cells in a TLR9-independent manner [20].

**Table 1.**
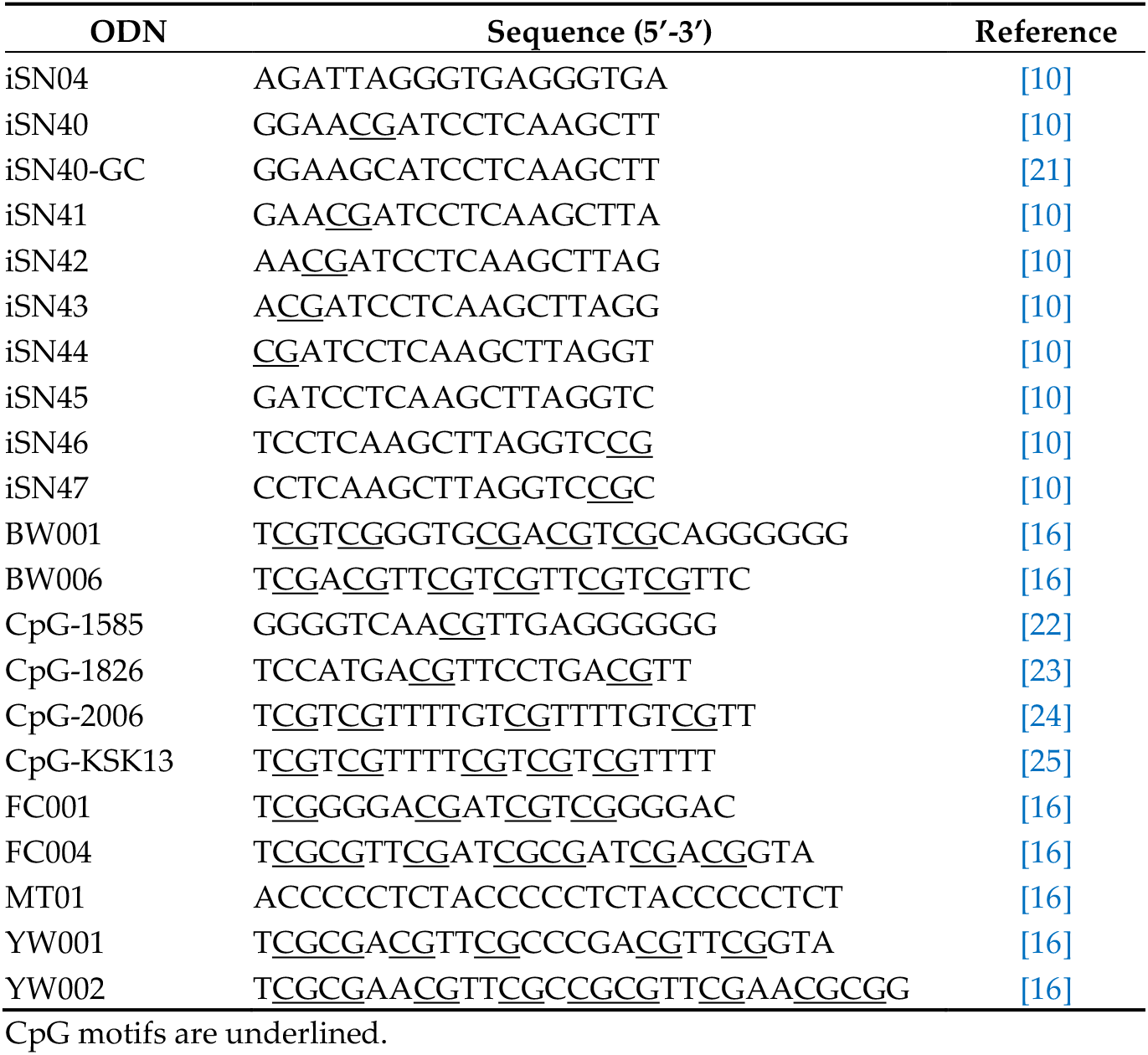
ODN sequences.

Furthermore, CpG-ODNs are known to inhibit the receptor activator of nuclear factor-κB ligand (RANKL)-induced osteoclastogenesis in a TLR9-dependent manner. CpG-1826 blocks osteoclast differentiation of murine bone marrow macrophages (mBMM) [23,26,27] but not of human peripheral blood monocytes (hPBMC) [23]. On the other hand,CpG-2006 works on hPBMC but not on mBMM. CpG-KSK13 was therefore designed to inhibit osteoclastogenesis in both mBMM and hPBMC [23]. CpG-1585 prevents osteoclast formation of the murine monocyte/macrophage cell line RAW264.7 by upregulating A20 deubiquitinase [22]. Similarly, YW001, YW002, and FC004, which were originally identified as CpG-osteoDNs [19], inhibit osteoclastic differentiation of RAW264.7 cells, whereas a non-CpG-osteoDN, MT01, does not affect osteoclastogenesis [28]. TLR9 dependencies of the anti-osteoclastogenic effects of these osteoDNs have not been clarified.

These studies demonstrated that CpG-ODNs, which regulate the differentiation of osteoblasts and osteoclasts, may be the medical seeds for osteoporosis therapy by elucidating their TLR9 dependencies. We have recently identified iSN40, a novel CpG-oste-oDN derived from the lactic acid bacterium genome, which strongly promotes the differentiation and calcification of MC3T3-E1 cells in a TLR9-independent manner [21]. To establish iSN40 as an anti-osteoporotic molecule, this study investigated the anti-osteoclastogenic effect and TLR9 dependency of iSN40 on RAW264.7 cells.

## 2. Materials and Methods

### 2.2. Chemicals

The ODNs used in this study (Table 1), in which all phosphodiester bonds were phos-phorothioated (PS) to enhance nuclease resistance, were synthesized and HPLC-purified (GeneDesign, Osaka, Japan). 6-FAM-ODNs are PS-ODNs conjugated to 6-carboxyfluo-recein at their 5’ ends (GeneDesign). All ODNs were dissolved in endotoxin-free water. Recombinant murine RANKL (ab129136; Abcam, Cambridge, UK) was dissolved in phos-phate-buffered saline (PBS). A TLR9 inhibitor, dihydrochloride (E6446) (MedChem Express, Monmouth Junction, NJ, USA), was dissolved in dimethyl sulfoxide. Ascorbic acid (Fujifilm Wako Chemicals, Osaka, Japan) was dissolved in endotoxin-free water. Equal volumes of solvents were used as the negative controls.

### 2.2. Culture and Osteoclastogenesis of RAW264.7 Cells

All cells were cultured at 37 °C with 5% CO_2_ throughout the experiments. RAW264.7 cells (91062702; Public Health England, London, UK) were maintained in DMEM (Nacalai, Osaka, Japan) with 10% fetal bovine serum (FBS) (HyClone; GE Healthcare, Salt Lake City, UT, USA) and a mixture of 100 units/ml penicillin and 100 μg/ml streptomycin (P/S) (Nacalai). For osteoclastogenesis, RAW264.7 cells were seeded on fresh dishes or plates, and 30 ng/ml RANKL was added the next day (defined as day 0). ODNs were simultane-ously treated with RANKL. To inhibit the TLR9 function, 1 μM E6446 was administered 3 h before ODN treatment. The medium containing RANKL, ODN, and E6446 was replaced every 2-3 days [29].

### 2.3. Tartrate-Resistant Acid Phosphatase (TRAP) Staining

RAW264.7 cells were seeded on 24-well plates (1.0 × 10^4^ cells/well) and induced oste-oclastogenesis in the RANKL-containing medium for 5-7 days. The TRAP enzymatic activity of the differentiated osteoclasts was visualized using TRAP Staining Kit (Fujifilm Wako Chemicals) according to the manufacturer’s instructions. Cell nuclei were stained with 1% methylene blue in 3% acetic acid. Bright-field images were captured using EVOS FL Auto microscope (AMAFD1000; Thermo Fisher Scientific, Waltham, MA, USA). The number of nuclei in TRAP^+^ multinucleated osteoclasts (with >3 nuclei) per field was counted using Image J software version 1.52u (National Institute of Health, Bethesda, MD, USA) [30].

### 2.4. Quantitative Real-Time RT-PCR (qPCR)

RAW264.7 cells were seeded on 30-mm (2.0 × 10^4^ cells/dish) or 60-mm (1.0 × 10^5^ cells/dish) dishes and induced osteoclastogenesis for 1-5 days. Total RNA from the cells was isolated using NucleoSpin RNA Plus (Macherey-Nagel, Düren, Germany) and was reverse transcribed using ReverTra Ace qPCR RT Master Mix (TOYOBO, Osaka, Japan). qPCR was performed using GoTaq qPCR Master Mix (Promega, Madison, WI, USA) with the StepOne Real-Time PCR System (Thermo Fisher Scientific). The amount of each transcript was normalized to that of tyrosine 3-monooxygenase/tryptophan 5-monooxygenase activation protein zeta (*Ywhaz*) gene. Results are expressed as fold-changes. Primer sequences are listed in Table 2.

**Table 2.**
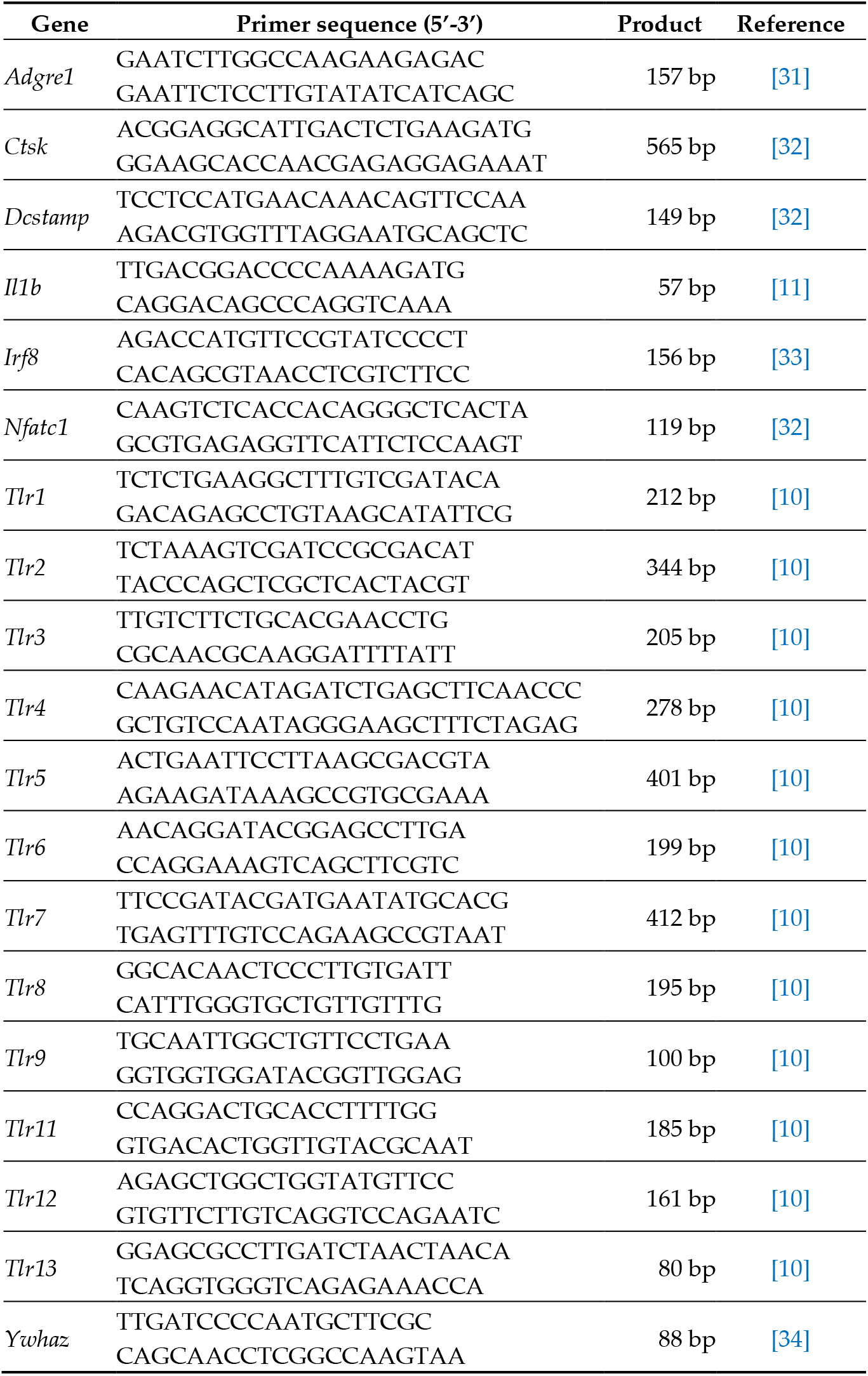
Primer sequences for qPCR.

### 2.5. ODN Incorporation Assay

RAW264.7 cells were seeded on 8-well glass chamber slides (3,000 cells/well) (SCS-N08; Matsunami Glass, Osaka, Japan) and treated with 5 μg/ml 6-FAM-iSN40 or 6-FAM-iSN40-GC for 30-60 min the next day. The cells were then washed with PBS, fixed with 2% paraformaldehyde, and stained with DAPI (Nacalai). Fluorescence images were captured using EVOS FL Auto microscope [10,11,15].

### 2.6. Multicanonical Molecular Dynamics (McMD) Simulation

The conformations of iSN40, iSN40-GC, and iSN41 were simulated by trivial-trajectory parallelization (TTP)-McMD method [35] as described previously [10,21,36]. Briefly, the initial structure of an ODN molecule was built as a DNA helix model by NAB in AmberTools [37] and the force field of amber ff99bsc0 was applied [38]. TTP-McMD was conducted to sample the equilibrated conformations at 310 K in the implicit solvent of water (igb = 1). The energy range of the multicanonical ensemble covered 270 K to 400 K. Sixty trajectories were used and 5 iterative runs were performed to achieve a flat distribution along the energy range. The production run was conducted for 100 ns in each trajectory (total 6 μs). Snapshots were saved every 200 ps. A total of 30,000 snapshots were sampled, and the 7033, 4039, and 5823 conformations of iSN40, iSN40-GC, and iSN41 at 310 K were obtained by the reweighting method. The representative ODN structures were taken from the centroid of structural clustering using The AmberTools22 [39]. Structure images were generated by UCSF Chimera [40].

### 2.7. Principal Component Analysis (PCA)

The molecular structures of iSN40, iSN40-GC, and iSN41 obtained by TTP-McMD were subjected to PCA. All sampled conformations of the three ODNs were used to construct principal components (PCs). The sines and cosines of 122 dihedral angles in the DNA-backbone conformation were used as the elements of the feature vector per conformation. The vector contained 244 dimensions. Variance-covariance matrices (244 × 244 dimensions) were calculated from the vectors of the sampled conformations. The matrices were diagonalized, and the eigenvalues and eigenvectors were obtained. The projection of the largest eigenvalue onto the eigenvector corresponded to the first and second components of the PCA (PC1 and PC2). These calculations were performed using Python scripts [14].

### 2.8. Coculture of RAW264.7 and MC373-E1 Cells

MC3T3-E1 cells (RCB1126; RIKEN BioResource Research Center, Tsukuba, Japan) were maintained in coculture medium consisting of α-MEM (Nacalai) with 10% FBS and P/S. For coculture, RAW264.7 cells (3.0 × 10^4^ cells/well) and MC3T3-E1 cells (1.0 × 10^4^ cells/well) were simultaneously seeded in coculture medium on 24-well plates [41,42]. The next day (defined as day 0), the cells were treated with 100 ng/ml RANKL, 50 μg/ml ascorbic acid, and ODNs to induce osteoclastogenesis.

### 2.9. Statistical Analysis

Results are presented as mean ± standard error. Statistical comparison among multiple groups was performed using Tukey-Kramer test after one-way analysis of variance. Statistical significance was set at *p* < 0.05.

## 3. Results

### 3.1. iSN40 Inhibits RANKL-Induced Osteoclastogenesis of RAW264.7 Cells

RAW264.7 cells were induced to osteoclastogenic differentiation using RANKL, treated with iSN40, and then stained for TRAP enzymatic activity, a marker of mature osteoclasts. As shown in Figure 1A, at day 7 after differentiation, multinucleated TRAP^+^ osteoclasts were formed in the RANKL-treated control group, but their osteoclastogenesis was interfered by iSN40. The effect of iSN40 was concentration-dependent. Osteoclast formation was significantly but incompletely suppressed by iSN40 at 0.1 μM, absolutely inhibited at 0.3 and 1.0 μM, and partially downregulated at 3.0 μM (Figure 1B, displayed logarithmically). The results indicated that iSN40 inhibits RANKL-induced osteoclastogenesis of RAW264.7 cells at optimal doses of 0.3–1.0 μM, which concentration range overlaps with that of other anti-osteoclastogenic CpG-ODNs [22,23,25–28]. In the following experiments, RAW264.7 cells were treated with 0.3 μM ODNs unless otherwise noted.

**Figure 1.**
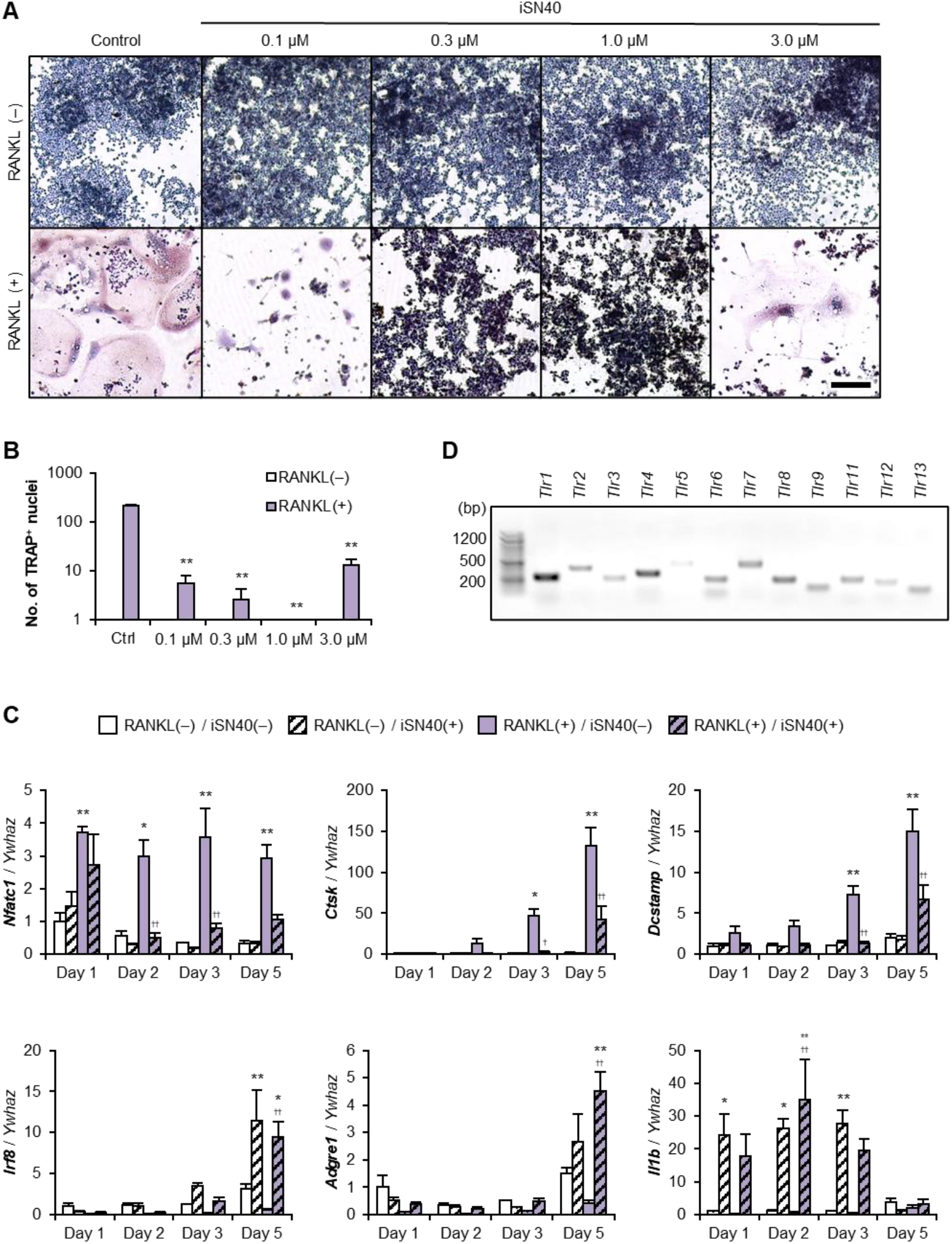
iSN40 inhibits RANKL-induced osteoclastogenesis of RAW264.7 cells. (**A**) Representative images of TRAP staining of RAW264.7 cells treated with 30 ng/ml RANKL and 0.1–3.0 μM iSN40 for 7 days. Scale bar, 100 μm. (**B**) The number of nuclei in TRAP^+^ osteoclasts was quantified. No TRAP^+^ cells were observed in any of the RANKL(-) groups. ** *p* < 0.05 vs RANKL(+)/control. *n* = 4 fields. (**C**) qPCR results of RAW264.7 cells treated with 30 ng/ml RANKL and 0.3 μM iSN40. \**p* < 0.05, ** *p* < 0.01 vs RANKL(-)/iSN40(-); ^†^ *p* < 0.05, ^††^ *p* < 0.01 vs RANKL(+)/iSN40(-) on each day. *n* = (**D**) Total RNA from RAW264.7 cells was subjected to RT-PCR (40 cycle), and then the PCR products of TLR genes were subjected to 1.5% agarose gel electrophoresis and stained with ethidium bromide.

qPCR revealed the iSN40-dependent gene expression changes in RAW264.7 cells (Figure 1C). iSN40 significantly downregulated RANKL-induced osteoclastogenic gene expression; nuclear factor of activated T cell 1 (*Nfatc1*) is the RANKL-responsive transcription factor that initiates osteoclast differentiation [11], cathepsin K (*Ctsk*) is an osteoclastspecific cysteine protease for bone resorption, and dendrocyte expressed seven transmembrane protein (*Dcstamp*) is the essential for cell fusion to form multinucleated osteoclasts. On the other hand, iSN40 significantly upregulated anti-osteoclastogenic transcription even in the presence of RANKL; interferon regulatory factor 8 (*Irf8*) is the competing transcription factor for NFATc1, which enhances macrophagic and suppresses osteoclastogenic gene expression [43]. Consistent with this, iSN40 strikingly induced a macrophage marker, adhesion G protein-coupled receptor E1 (*Adgre1*; F4/80), despite RNAKL treatment. iSN40 significantly increased interleukin 1β (IL-1β) (*Il1b*) mRNA levels independently of RANKL. Since RAW264.7 cells expressed all TLR genes (Figure 1D), it was hypothesized that iSN40 serves as a CpG-ODN recognized by TLR9 to induce immune responses.

### 3.2. iSN40 Inhibits Osteoclastogenesis in a TLR9-Dependent Manner

The TLR9 dependency of iSN40 was investigated using a TLR9 inhibitor, E6446, and a known CpG-ODN, CpG-2006, which has been reported to activate TLR9 in RAW264.7 cells [24]. As shown Figure 2A, CpG-2006 completely inhibited RANKL-induced osteoclastogenesis of RAW264.7 cells to the same extent as iSN40. As expected, pre-treatment with E6446 reversed the anti-osteoclastogenic effects of iSN40 and CpG-2006. Although TRAP^+^ cells were still small in the iSN40 and CpG-2006 groups with E6446, the reduction in the number of nuclei in TRAP^+^ cells by iSN40 or CpG-2006 was fully recovered by E6446 (Figure 2B). qPCR also showed that E6446 completely blocked iSN40- and CpG-2006-induced *Il1b* expression (Figure 2C). These data demonstrate that recognition of iSN40 by TLR9 is an essential step in the interruption of RANKL-induced osteoclast differentiation.

**Figure 2.**
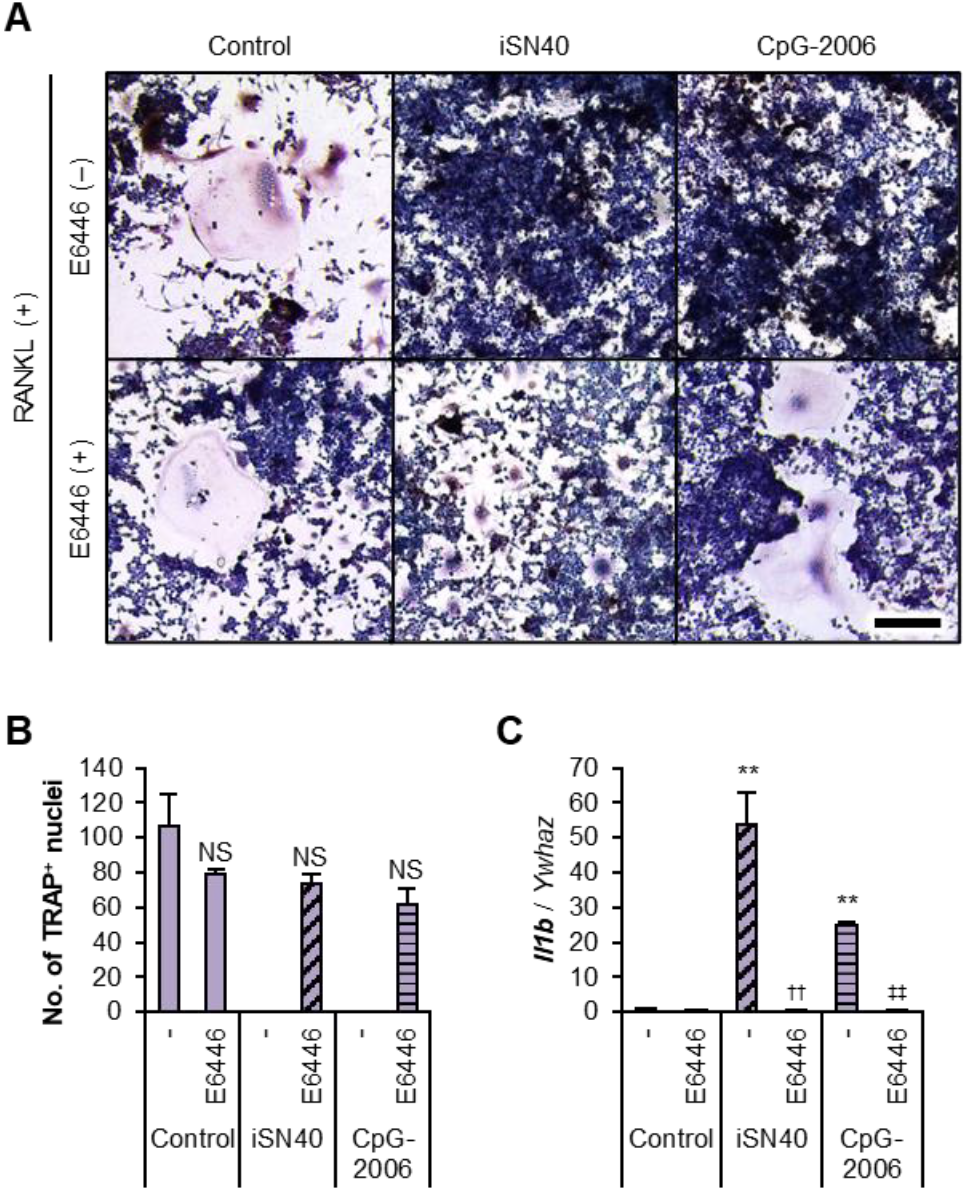
iSN40 inhibits osteoclastogenesis in a TLR9-dependent manner. (**A**) Representative images of TRAP staining of RAW264.7 cells treated with 30 ng/ml RANKL, 0.3 μM ODN, and 1 μM E6446 for 6 days. Scale bar, 100 μm. (**B**) The number of nuclei in TRAP^+^ osteoclasts was quantified. No TRAP^+^ cells were observed in the iSN40/E6446(-) and CpG-2006/E6446(-) groups. NS, no significant difference vs control/E6446(-). *n*= 4 fields. (**C**) qPCR results of RAW264.7 cells treated with 30 ng/ml RANKL, 0.3 μM ODNs, and 1 μM E6446 for 24 h. ** *p* < 0.01 vs control/E6446(-), ^††^ *p* < 0.01 vs iSN40/E6446(-), ^‡‡^ *p* < 0.01 vs CpG-2006/E6446(-). *n* = 3.

### 3.3. The CpG Motif Within iSN40 Is Essential for Its Anti-Osteoclastogenic Effect

TLR9 recognizes the unmethylated CpG di-deoxynucleotide motifs within CpG-ODNs [44]. To investigate the impact of the CpG motif within iSN40 (5’-GGA A<u>CG</u> ATC CTC AAG CTT-3’) on osteoclastogenesis, iSN40-GC (5’-GGA A<u>GC</u> ATC CTC AAG CTT), in which the CG sequence was substituted with GC (underlined sequences), was used. iSN40-GC did not inhibit the RANKL-induced osteoclast formation of RAW264.7 cells at all (Figures 3A and 3B). Importantly, MT01, another osteoDN having no CpG motif, also showed no action on osteoclastogenesis, demonstrating that the pro-osteogenic and anti-osteoclastogenic effects of osteoDNs are independent and can be separated. qPCR reproducibly showed that iSN40-GC did not initiate immune responses such as *Il1b* expression and did not downregulate osteoclastogenic gene expression including *Nfatc1, Ctsk*, and *Dcstamp* (Figure 3C). These data clearly represent that the CpG motif is essential for the anti-osteoclastogenic effect of iSN40 via recognition by TLR9.

**Figure 3.**
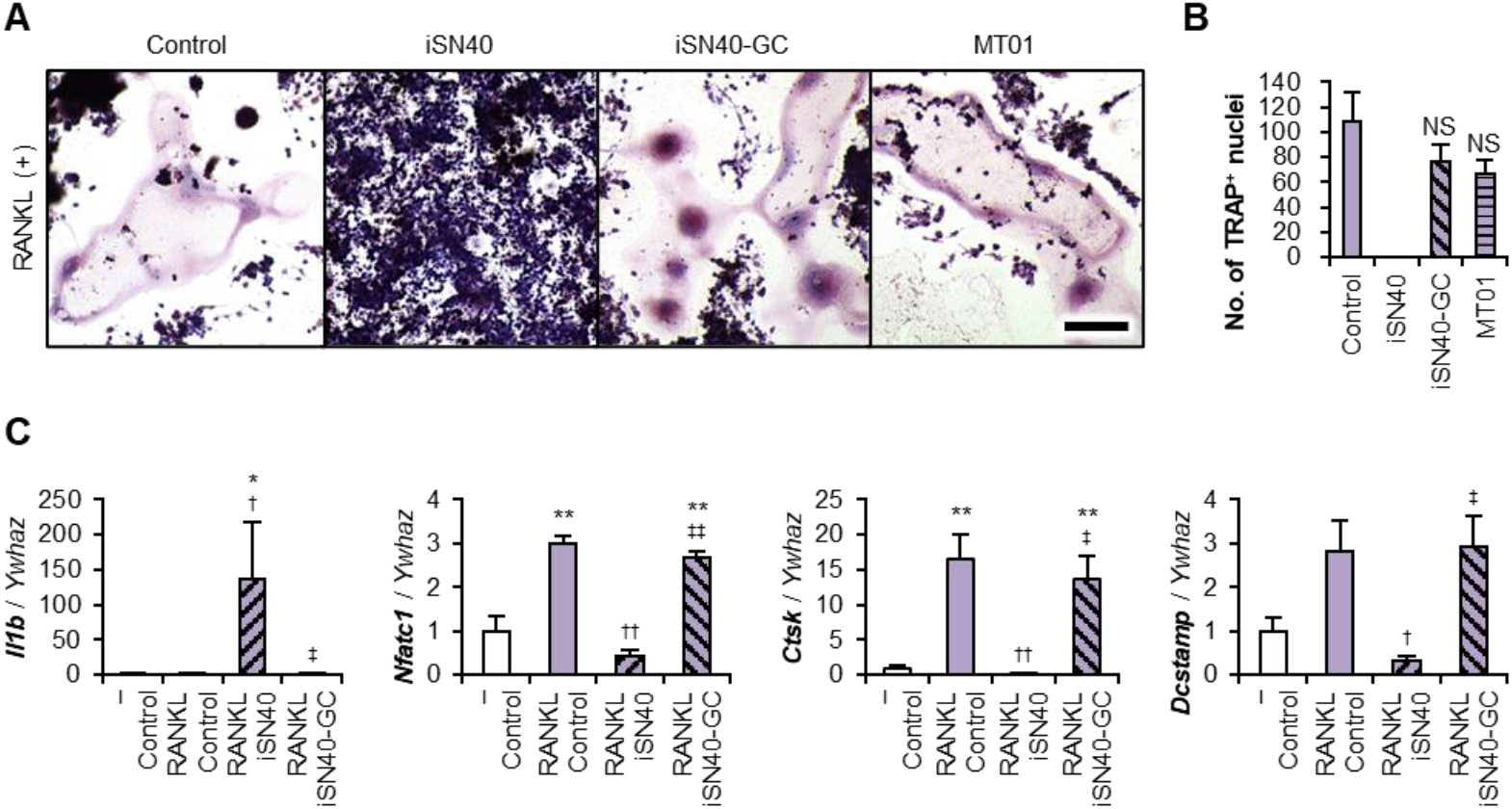
The CpG motif is essential for anti-osteoclastogenic effect of iSN40. (**A**) Representative images of TRAP staining of RAW264.7 cells treated with 30 ng/ml RANKL and 0.3 μM ODN for 7 days. Scale bar, 100 μm. (**B**) The number of nuclei in TRAP^+^ osteoclasts was quantified. No TRAP^+^ cells were observed in the iSN40 group. NS, no significant difference vs control. *n* = 4 fields. (**C**) qPCR results of RAW264.7 cells treated with 30 ng/ml RANKL, 0.3 μM ODNs for 24 h (*Il1b*) or 5 days (*Nfatc1, Ctsk*, and *Dcstamp*). * *p* < 0.05, ** *p* < 0.01 vs control/RANKL(-); ^†^ *p* < 0.05, ^††^ *p* < 0.01 vs control/RANKL(+); ^‡^ *p* < 0.05, ^‡‡^ *p* < 0.01 vs iSN40/RANKL(+). *n* = 3.

### 3.4. The CpG Motif of iSN40 Is Important for Intracellular Incorporation

CpG-ODNs must be taken up intracellularly to associate with TLR9 in endosomes [45]. The previous study has demonstrated that both iSN40 and iSN40-GC were similarly incorporated into the cytoplasm of MC3T3-E1 cells, which do not express TLR9, within 30 min without a carrier [21]. As shown in Figure 4, 6-FAM-iSN40 administered to RAW264.7 cells was autonomously internalized into the cytoplasm within 30 min and localized in endosome-like vesicles within 60 min. While 6-FAM-iSN40-GC was less incorporated compared to iSN40, and the vesicles were not clearly observed. These results suggest that the CpG motif of iSN40 is important for its intracellular incorporation and subcellular localization, probably to TLR9 in endosomes, which is related to the anti-osteoclastogenic effect.

**Figure 4.**
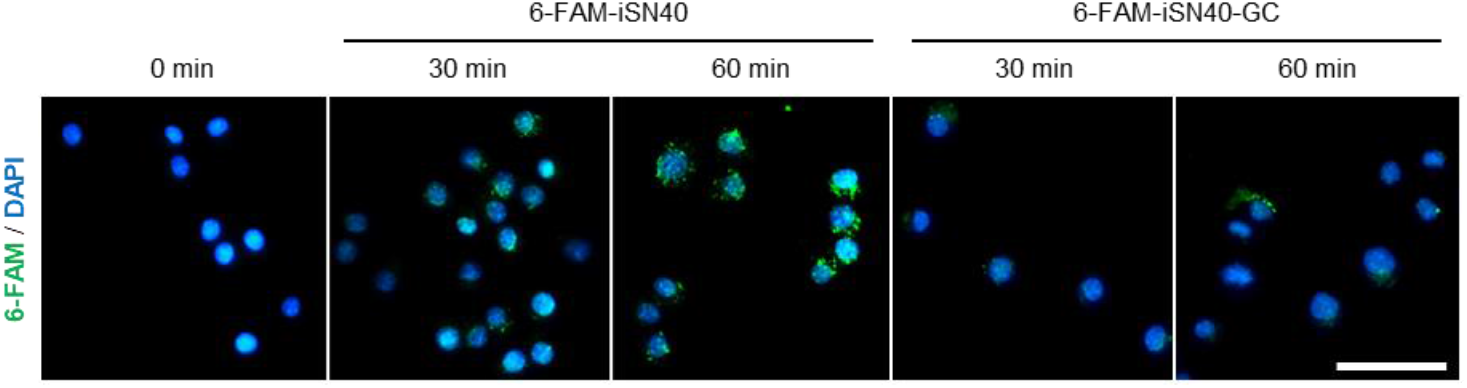
Intracellular incorporation of iSN40. Representative fluorescence images of RAW264.7 cells treated with 5 μg/ml 6-FAM-iSN40 or 6-FAM-iSN40-GC for 30-60 min. Scale bar, 50 μm.

### 3.5. The Location of the CpG Motif Is Essential for iSN40 Activity

It is not necessarily the case that all CpG-ODNs can be TLR9 ligands [44]. The iSN40 variants, iSN41–iSN47 (Table 1), have a CpG motif (except for iSN45, which has no CpG motif as a negative control) but are not osteoDNs [21]. Anti-osteoclastogenic effects and immune responses of these ODNs were investigated. iSN41 and iSN42 completely and iSN43 partially inhibited osteoclastogenesis, but the others showed no significant effects (Figures 5A and 5B). Correspondingly, iSN41 markedly induced *Il1b* expression as well as iSN40 (Figure 5C). iSN41, the most homologous to iSN40, showed almost the same bioactivities as iSN40 in RAW264.7 cells. Overall, iSN40–iSN43 with a CpG motif in their 5’ regions (5’-CpG) exhibited anti-osteoclastogenic and immunostimulatory effects, whereas iSN46 and iSN47 with the 3’-CpG dinucleotide did not. iSN44 with a CpG motif at its 5’ terminus and iSN45 with no motif also showed no activities. These results are consistent with the previous study reporting that CpG-ODNs with a 5’-CpG are more active than those with a 3’-CpG [46]. These suggest that the CpG motif located at the 5-6th residues is essentially required for the bioactivities of iSN40.

**Figure 5.**
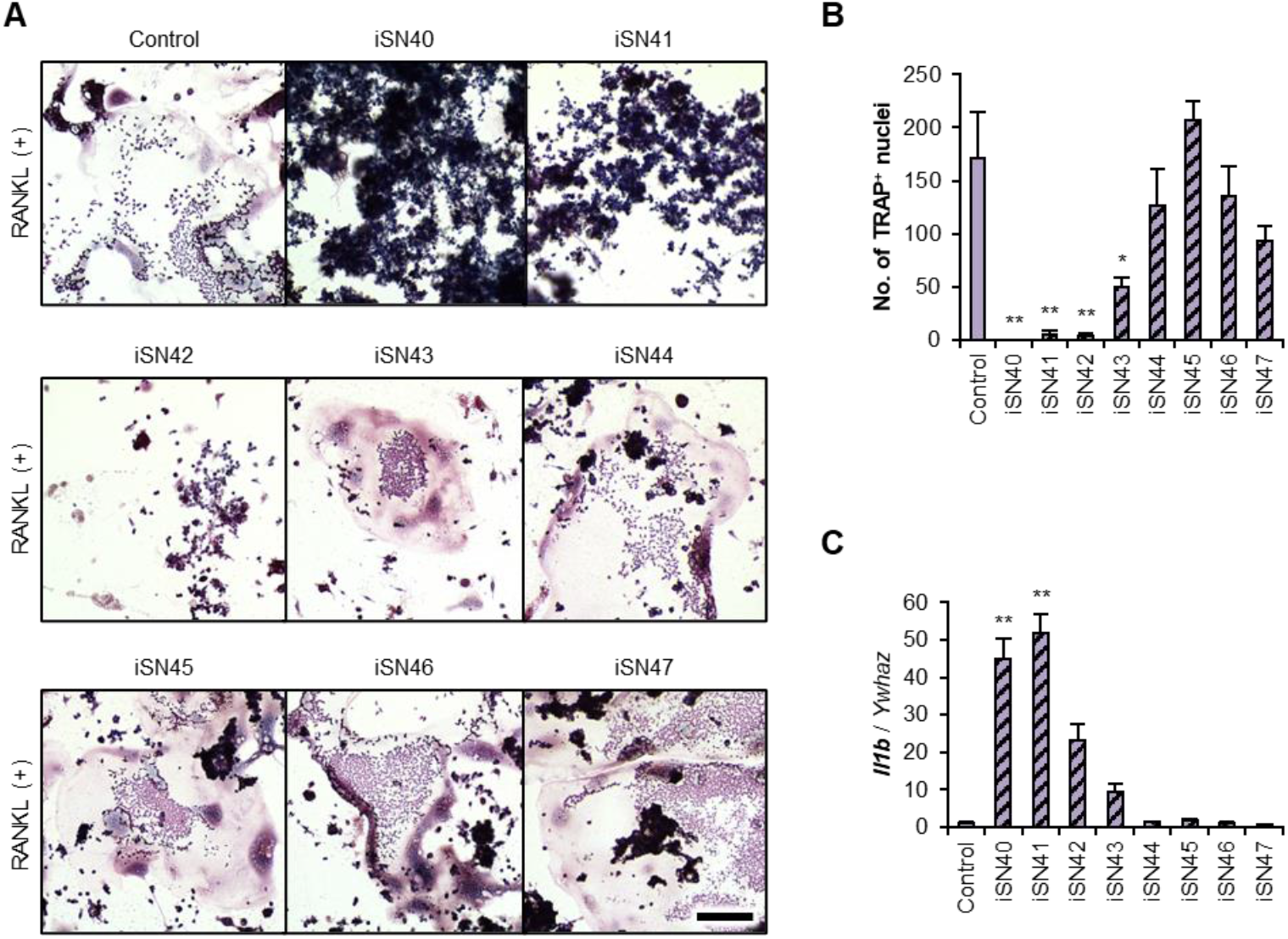
Anti-osteoclastogenic effects of the iSN40 variants. (**A**) Representative images of TRAP staining of RAW264.7 cells treated with 30 ng/ml RANKL and 0.3 μM ODN for 7 days. Scale bar, 100 μm. (**B**) The number of nuclei in TRAP^+^ osteoclasts was quantified. No TRAP^+^ cells were observed in the iSN40 group. * *p* < 0.05, ** *p* < 0.01 vs control. *n* = 4 fields. (**C**) qPCR results of RAW264.7 cells treated with 30 ng/ml RANKL and 0.3 μM ODN for 24 h. * *p* < 0.05, ** *p* < 0.01 vs control. *n* = 3.

### 3.6. Molecular Simulation of the iSN40 Structure

iSN40 and iSN41 exerted anti-osteoclastogenic effects in a TLR9-dependent manner, but a non-CpG-ODN, iSN40-GC, did not. On the other hand, iSN40 and iSN40-GC can promote osteogenesis independent of TLR9, but iSN41 does not [21]. These results suggest that the pro-osteogenic and anti-osteoclastogenic effects of iSN40 are based on different mechanisms and can be functionally separated. To analyze the relationship between molecular structures and functions of iSN40, iSN40-GC, and iSN41, their conformations were computationally simulated using TTP-McMD. The resulting structures were subjected to PCA analysis and clustered to abstract their representative conformations (Figure 6A).

**Figure 6.**
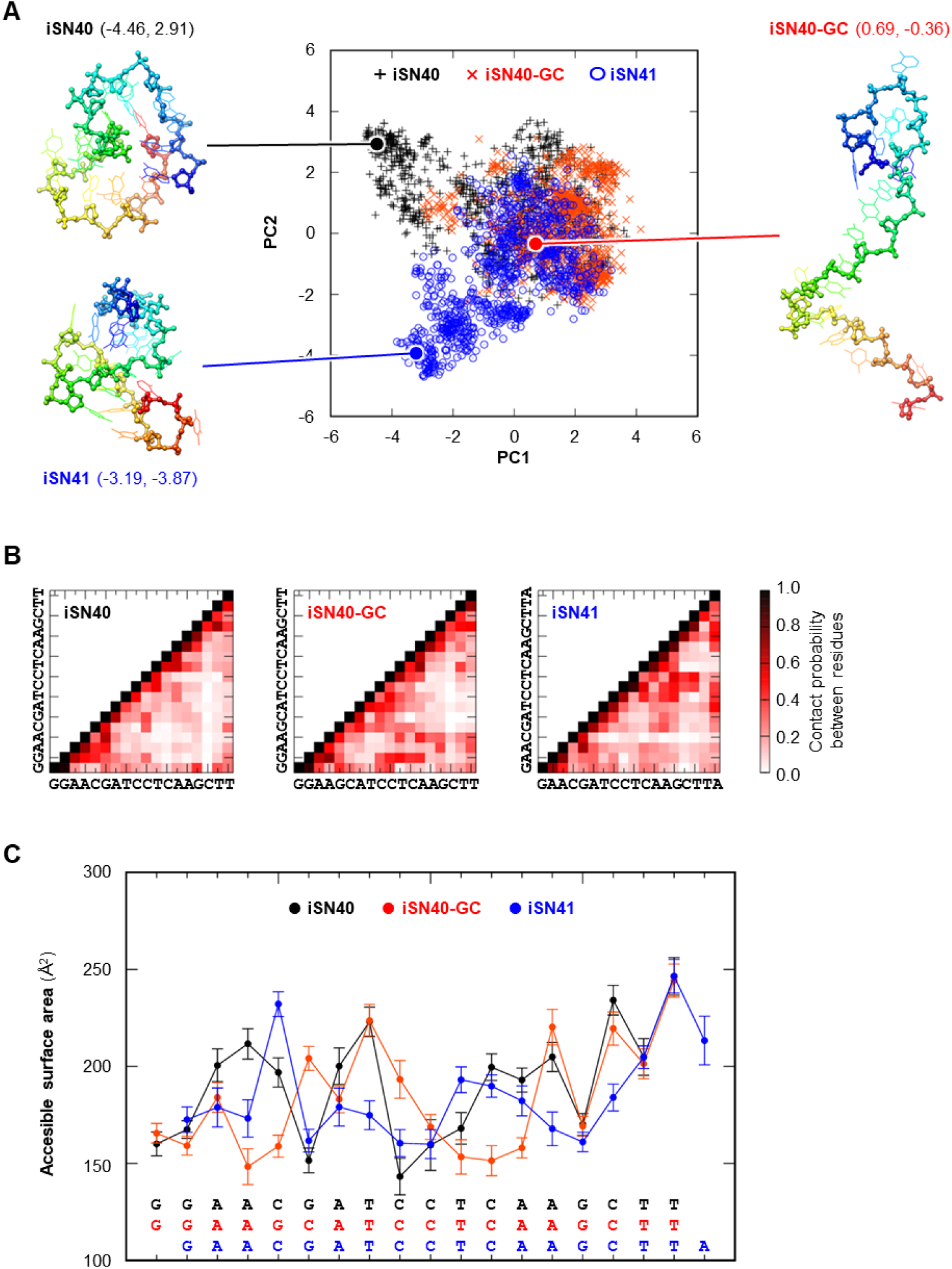
Molecular simulation of the structures of iSN40, iSN40-GC, and iSN41. (**A**) The structures of iSN40, iSN40-GC, and iSN41 simulated by TTP-McMD are shown in a two-dimensional plot on the PCA space reconstructed by PC1 (contribution, 0.083) and PC2 (0.064). The molecular structures, shown as a rainbow-colored stick model from the 5’ end (blue) to the 3’ end (red), are representative conformations taken from the centroid of the structural clustering. The PCA coordinate (PC1, PC2) of each conformation is indicated. (**B**) The contact probabilities between residues within iSN40, iSN40-GC, and iSN41. (**C**) The accessible surface area of the residues of iSN40, iSN40-GC, and iSN41.

The molecular structures of iSN40, iSN40-GC, and iSN41 partially overlapped, probably because their sequences were nearly identical. However, detailed analysis revealed sub-moleculer differences. As shown in Figure 6B, the contact probabilities (analogous to distance) between residues within iSN40 were similar to those of iSN40-GC. In particular, the 3’ terminal sequences, AGCTT, were structurally open (residues not contacting each other) in iSN40 and iSN40-GC, whereas in iSN41, the 15-16th CT interacted with the 9-10th CT. Also, the accessible surface area (analogous to exposure) patterns of AGCTT were closely resembled in iSN40 and iSN40-GC, but not in iSN41 (Figure 6C). These data suggest that the common AGCTT properties of iSN40 and iSN40-GC may contribute to their TLR-independent pro-osteogenic effects. Focusing on the CpG motifs of iSN40 and iSN41, the 6th G was in close contact with the 4-5th AC in iSN40 (contact between the 5th G and the 3-4th AC in iSN41), suggesting to be the basis of their TLR9-dependent anti-osteoclastogenic actions.

### 3.7. iSN40 Exerts Anti-Osteoclastogenic Effect in the Presence of Osteoblasts

Finally, the anti-osteoclastogenic effect of iSN40 was tested in the presence of osteoblasts by coculturing RAW264.7 cells and MC3T3-E1 osteoblasts [41,42], because osteoclasts and osteoblasts mutually regulate their differentiation through RANKL-RANK signaling to maintain the balance between bone resorption and formation in vivo [1]. Since both iSN40 and iSN40-GC promote osteoblast differentiation independently of TLR9 [21], their pro-osteogenic effects may indirectly influence osteoclastogenesis of RAW264.7 cells in coculture with MC3T3-E1 cells. Confirmation of the anti-osteoclastogenic effect of iSN40 in the presence of osteoblasts is therefore important for animal studies and clinical application. The cocultured RAW264.7 and MC3T3-E1 cells were subjected to RANKL-induced osteoclastogenesis for 4 days. As shown in Figure 7A, undifferentiated RAW264.7 cells, differentiated TRAP^+^ multinucleated osteoclasts, and fibroblast-like MC3T3-E1 osteoblasts were in close contact with each other, allowing paracrine signaling among them during differentiation in this system.

**Figure 7.**
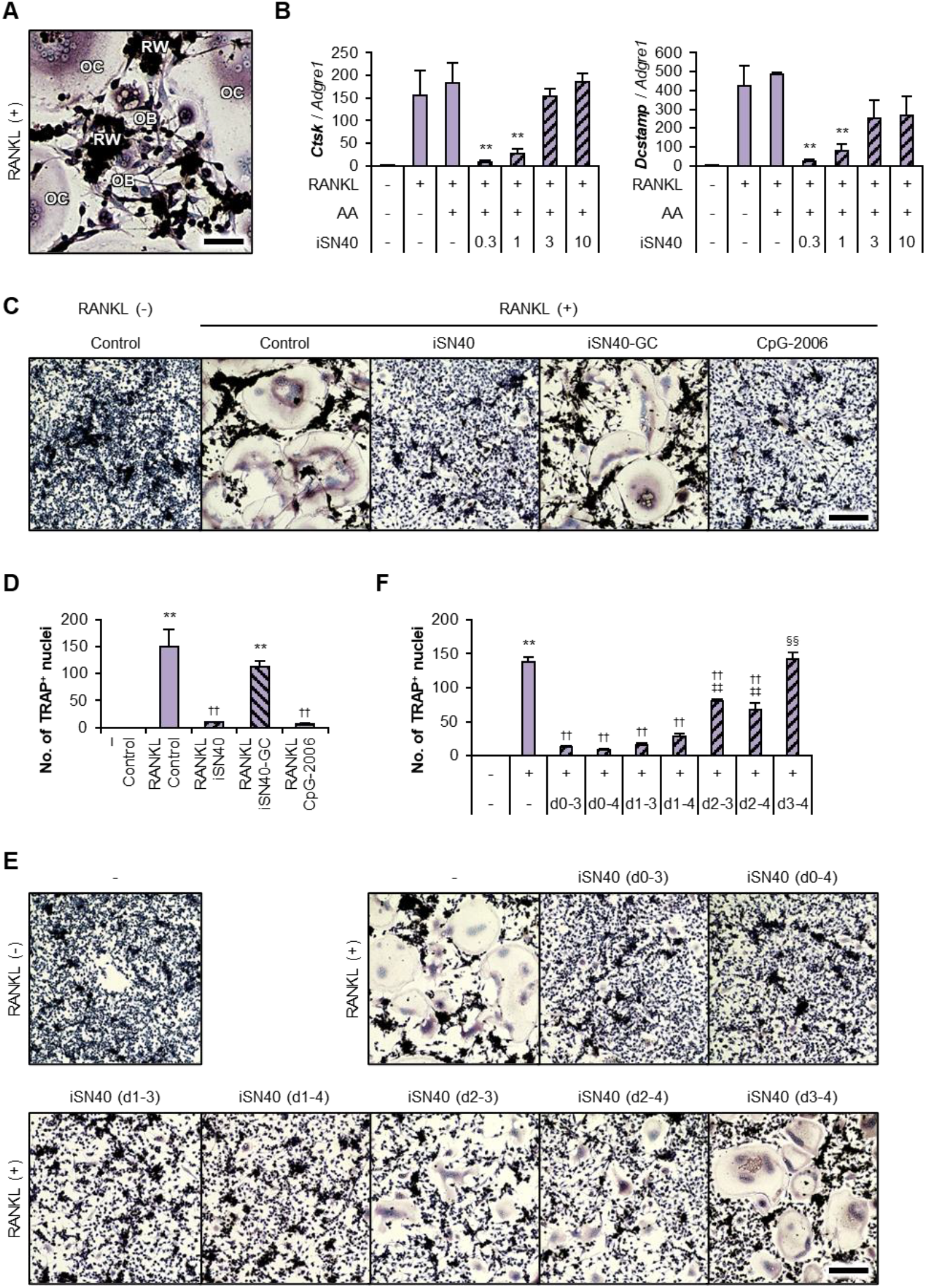
Anti-osteoclastogenic effect of iSN40 in the coculture of RAW264.7 and MC3T3-E1 cells. (**A**) Representative images of TRAP staining of cocultured RAW264.7 and MC3T3-E1 cells treated with 100 ng/ml RANKL for 4 days. RW, undifferentiated RAW264.7 cells (dense nuclei in black); OC, differentiated TRAP^+^ multinucleated osteoclasts; OB, fibroblast-like MC3T3-E1 osteoblasts. Scale bar, 25 μm. (**B**) qPCR results of cocultured RAW264.7 and MC3T3-E1 cells treated with 100 ng/ml RANKL, 50 μg/ml ascorbic acid (AA), and 0.3–10 μM iSN40 for 4 days. ** *p* < 0.01 vs control/RANKL(+)/AA(+). *n* = 3. (**C**) Representative images of TRAP staining of cocultured RAW264.7 and MC3T3-E1 cells treated with 100 ng/ml RANKL and 0.3 μM ODN for 4 days. Scale bar, 100 μm. (**D**) The number of nuclei in TRAP^+^ osteoclasts was quantified. ** *p* < 0.01 vs control/RANKL(-), ^††^ *p* < 0.01 vs control/RANKL(+). *n* = 3 fields. (**E**) Representative images of TRAP staining of cocultured RAW264.7 and MC3T3-E1 cells treated with 100 ng/ml RANKL for 4 days and 0.3 μM iSN40 for defined periods. d, differentiation days with iSN40 treatment. Scale bar, 100 μm. (**F**) The number of nuclei in TRAP^+^ osteoclasts was quantified. ** *p* < 0.01 vs control/RANKL(-); ^††^ *p* < 0.01 vs control/RANKL(+); ^‡‡^ *p* < 0.01 vs d0-3, d0-4, d1-3, and d1-4; ^§§^ *p* < 0.01 vs d2-3 and d2-4. *n* = 3 fields.

Ascorbic acid has been widely used not only to induce differentiation of MC3T3-E1 cells [47], but is also known to inhibit RANKL-induced osteoclastogenesis of RAW264.7 cells at higher doses [48], which is promising as a potential nutritional vitamin for osteoporosis therapy [49]. The RANKL-induced expression levels of the osteoclastogenic genes, *Ctsk* and *Dcstamp*, were quantified in the coculture condition treated with RANKL and 50 μg/ml ascorbic acid. Transcripts were normalized to that of *Adgre1*, which is expressed only in RAW264.7 cells, not in MC3T3-E1 cells. As shown in Figure 7B, ascorbic acid did not alter the RANKL-induced *Ctsk* and *Dcstamp* levels in this system. iSN04 significantly inhibited osteoclastogenesis at the dose of 0.3-1.0 μM but not at 3–10 μM, as observed in the single RAW264.7 cell culture (Figures 1A and 1B). TRAP staining clearly visualized that iSN40 and CpG-2006, but not iSN40-GC, inhibited the RANKL-induced osteoclastogenesis in the coculture system (Figures 7C and 7D). These results demonstrate that iSN40 can exert its anti-osteoclastogenic effect based on its CpG motif and TLR9 even in the presence of osteoblasts.

In the above coculture experiments, iSN40 was treated throughout the 4 days of RANKL-induced differentiation. To determine the effective period of anti-osteoclastogenic treatment, iSN40 was administered and removed at different time points. As shown in Figures 7E and 7F, iSN40 treatment during differentiation days 1 to 3 (d1-3) was sufficient to inhibit osteoclast formation compared to the conventional full treatment (d0-4). While iSN04 administration from day 2 (d2-3 and d2-4) partially allowed osteoclastogenesis, and that from day 3 (d3-4) showed no effect. These data suggest that the first 2 days of osteoclast differentiation are essential for the anti-osteoclastogenic mechanism of iSN40.

## 3. Discussion

This study demonstrated that a CpG-osteoDN, iSN40, completely inhibits RANKL-induced osteoclastic differentiation of RAW264.7 cells in a TLR9-dependent manner. The anti-osteoclastogenic effect of iSN40 was abolished by a TLR9 inhibitor or by substitution of its CpG motif with a GC sequence. iSN40 initiated immune responses such as IL-1β induction, resulting in abrogation of RANKL-induced osteoclastogenic gene expression. The optimal dose of iSN40 was 0.3–1.0 μM, which corresponds to that of known CpG-ODNs for TLR9-dependent immunostimulatory activities [50]. Structure simulation visualized that the CpG motif of iSN40 is exposed on the molecular surface, which is readily sensed by TLR9. Both of an iSN40 homolog, iSN41, and a different type of CpG-ODN, CpG-2006, inhibited osteoclastogenesis to the same extent as iSN40, indicating that the anti-osteoclastogenic functions of these ODNs are mediated by a common mechanism. These results clearly presented that the CpG motif within iSN40 was recognized by TLR9 and was essential to interfere with osteoclast formation of RAW264.7 cells. Immunostimulatory CpG-ODNs have been used as vaccine adjuvants and anti-cancer agents [9,51], and their safety has also been evaluated and confirmed. It suggests that iSN40 as a CpG-ODN would be safe when used to suppress bone resorption.

On the other hand, iSN40 was originally identified as an osteoDN that promotes osteoblast differentiation independent of TLR9 at the optimal dose of 1–10 μM [21]. iSN40 is expected to have a dual function, pro-osteogenic and anti-osteoclastogenic effects, which simultaneously accelerate bone formation and suppress bone resorption in vivo. This study demonstrated that iSN40 can exert at least its anti-osteoclastogenic effect in the coexistence of osteoblasts and osteoclasts. The pro-osteogenic action of iSN40 in the coculture system should be investigated as a next step. Like romosozumab, a double action on both osteogenesis and osteoclastogenesis is beneficial for osteoporosis therapy [6,7]. iSN40 may be a promising nucleic acid drug for osteoporosis by modulating bone remodeling. Note that the pro-osteogenic and anti-osteoclastogenic effects of iSN40 have been investigated in the murine cell lines, MC3T3-E1 and RAW264.7, respectively. Recognition of CpG-ODNs by TLR9 can be species-specific [24]. For example, CpG-1826 blocks osteoclastogenesis in mouse cells [23,26,27] but not in human cells [25]. The effects of iSN40 on human osteoblasts and osteoclasts need to be investigated in future studies for clinical application.

Although iSN40-GC without CpG motif promoted osteogenesis of MC3T3-E1 cells in the previous study [21], it did not inhibit osteoclastogenesis of RAW264.7 cells in this study. In contrast, iSN41 with CpG motif did not affect osteoblasts [21], while it disrupted osteoclast formation. These facts proved that pro-osteogenic and anti-osteoclastogenic effects of ODNs can be separated (Figure 8). The pro-osteogenic mechanisms of iSN40 and iSN40-GC are TLR9-independent, but the details including the direct target(s) are still unknown [21]. Another osteoDN, MT01, is also capable of directing osteogenesis but does not have CpG motifs [16–19]. While CpG-2006 is a CpG-ODN but interferes with bone morphogenic protein-induced osteoblastic differentiation in a TLR9-independent manner [20]. Basically, osteoblasts do not express TLR9 [21,52]. These studies present that the ODN functions on osteoblasts are not based on TLR9, regardless of their CpG motifs. These ODNs would serve as nucleic acid aptamers that specifically interact with the target molecules (typically proteins) in a manner similar to antibodies [53]. We have reported that bacterial genome-derived ODNs can be aptamers and regulate cell fate [10], leading to the hypothesis that iSN40, derived from the lactic acid bacterium genome, functions as an aptamer in osteoblasts. As computationally simulated, iSN40 forms a stable globular conformation (average radius, 1.01 nm) [21], which is a typical feature of a type of aptamer. If iSN40 is an aptamer, then iSN40-GC is also considered to be an aptamer targeting an identical molecule. Molecular simulation showed that the contact probabilities and the accessible surface area of the 3’ terminal sequence, AGCTT, are similar between iSN40 and iSN40-GC, but not iSN41, suggesting that this region may be the core structure for their aptamer function. Identification of the target molecule of iSN40 and iSN40-GC in osteoblasts is an important issue for their development as nucleic acid drugs for bone diseases.

**Figure 8.**
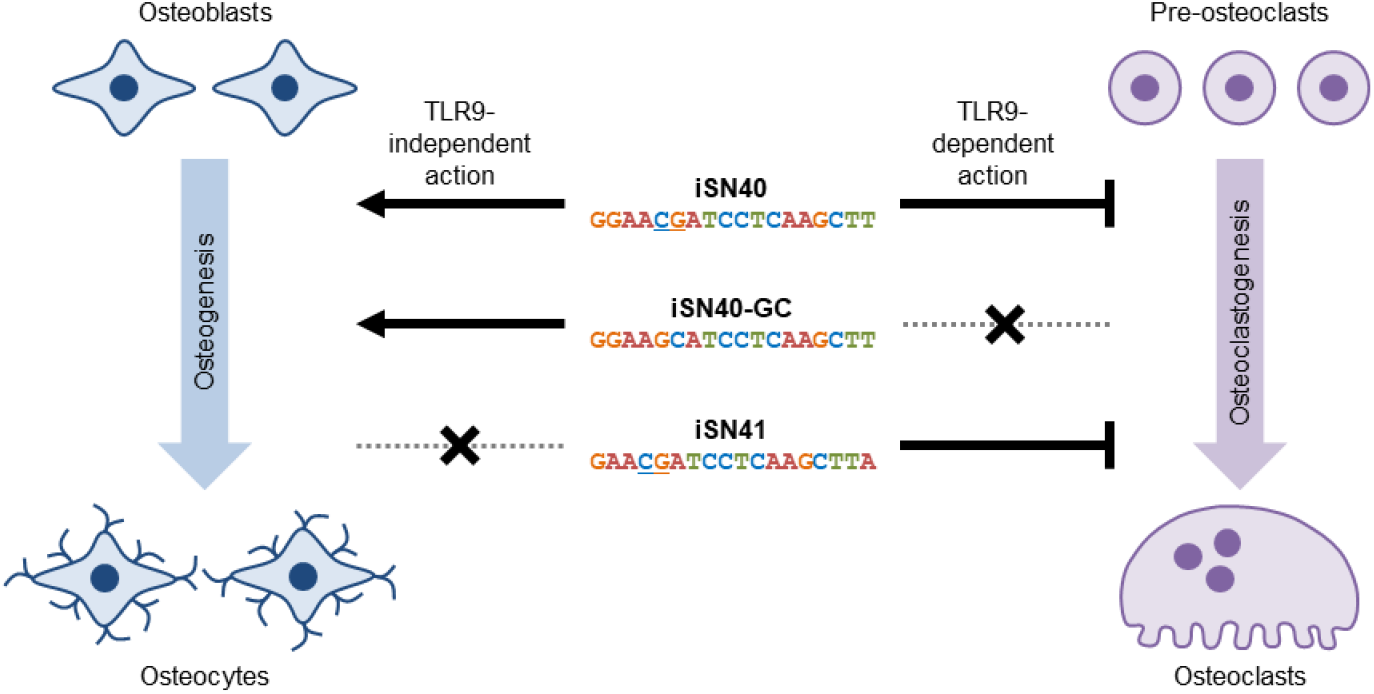
TLR9-independent pro-osteogenic and TLR9-dependent anti-osteoclastogenic effects of iSN40, iSN40-GC, and iSN41. CpG motifs are underlined.

Our studies successfully developed three ODNs; iSN40 with both pro-osteogenic and anti-osteoclastogenic effects, iSN40-GC which induces only osteogenesis, and iSN41 which inhibits only osteoclastogenesis (Figure 8). The combination or proper use of these ODNs will achieve fine control of bone remodeling to balance bone formation and resorption, ultimately opening the door to the treatment of osteoporosis with nucleic acid drugs. Osteoporosis affects approximately 200 million people in the world and is associated with to 8.9 million fractures per year [54]. Anti-sclerostin antibody, romosozumab [6,7], and anti-RANKL antibody, denosumab [55,56], are now widely used for osteoporosis therapy, however, antibody drugs are still expensive for a large number of patients worldwide. ODNs can be synthesized chemically, rapidly, and economically on a large scale [8,57]. iSN40, which exerts pro-osteogenic and anti-osteoclastogenic effects, is a potent seed to be studied and developed as a nucleic acid drug for sustainable osteoporosis therapy.

## 5. Conclusions

This study demonstrated that iSN40, the 18-base CpG-ODN derived from the lactic acid bacterium genome, strongly inhibited RANKL-induced osteoclastogenesis through the recognition of its CpG motif by TLR9. The previous study reported that iSN40 promote osteogenesis. The pro-osteogenic and anti-osteoclastogenic functions of iSN40 may be useful to maintain the balance between bone formation and bone resorption. iSN40 is a potential nucleic acid drug for osteoporosis therapy by regulating bone remodeling.

## Author Contributions

Conceptualization, T.T.; investigation, R.I., C.K., Y.N. and K.U.; resources, T.S.; writing—original draft preparation, T.T.; funding acquisition, C.K. and T.T. All authors have read and agreed to the published version of the manuscript.

## Funding

This research was funded by the Japan Society for the Promotion of Science (19K05948 and 22K05554 to T.T.) and by the Fund of Nagano Prefecture to Promote Scientific Activity (NPS2024306 to C.K.).

## Institutional Review Board Statement

Not applicable.

## Informed Consent Statement

Not applicable.

## Data Availability Statement

The data presented in this study are available on request to the corresponding author.

## Conflicts of Interest

The authors declare no conflicts of interest.

## Notes

### Competing Interest Statement

The authors have declared no competing interest.

